# Prognostic and Therapeutic Implications of *BRAF* Mutations in Acute Myeloid Leukemia

**DOI:** 10.1101/2025.10.14.682328

**Authors:** Darren Lee, Yazan Abu-Shihab, Kate Plas, Deedra Nicolet, Krzysztof Mrozek, Mark J. Routbort, Keyur P. Patel, Christopher J. Walker, Jill Buss, Andrew R. Stiff, Andrea Laganson, Courtney D. DiNardo, Naval G. Daver, Tapan M. Kadia, Farhad Ravandi, Andrew J. Carroll, Jonathan E. Kolitz, Bayard L. Powell, William G. Blum, Maria R. Baer, Guido Marcucci, Geoffrey L. Uy, Wendy Stock, Richard M. Stone, L. Jeffrey Medeiros, John C. Byrd, James S. Blachly, Robert L. Bowman, Jeffrey W. Tyner, Sanam Loghavi, Ann-Kathrin Eisfeld, Linde A. Miles

## Abstract

Mutations in the RAS/MAPK signaling pathway are recurrent in acute myeloid leukemia (AML), primarily involving *NRAS* and *KRAS*. In contrast, mutations in the gene encoding an effector protein, BRAF, occur at relatively lower frequencies in AML and are associated with poor outcomes. To date, no comprehensive analysis has assessed the clinical and molecular characteristics of *BRAF-*mutated AML. In this study, we report the identification of canonical and non-canonical *BRAF* mutations in ∼1% of 5,779 consecutive clinically and molecularly fully-annotated AML patients treated at two major United States Cancer Centers (50/5779 AML patients: 21 newly diagnosed AML; 9 relapsed/refractory; 20 newly diagnosed secondary AML). We performed single-cell multiomic analysis on a subset of AML samples. *BRAF* mutations were enriched in myelodysplasia-related AML (AML-MR), and most mutations were located outside the V600 hotspot. Single-cell multiomic profiling delineated *BRAF* mutation class-specific patterns of co-mutations, clonality, and immunophenotypes. Notably, *BRAF* mutations and other signaling co-mutation(s) could be found in the same cell, a finding that significantly diverges from prior studies of RAS-mutant AML. In this cohort, *BRAF*-mutant AML patients had poor overall survival with currently available treatments, including venetoclax-based regimens. Drug sensitivity data suggest possible avenues for targeted treatment of *BRAF*-mutated AML.

**Statement of Significance:** Canonical and non-canonical *BRAF* mutations are enriched in AML-MR and associate with poor survival outcomes. Single-cell multiomic profiling revealed unique co-mutation patterns and immunophenotypes that highlight RAS pathway addiction and nominate *BRAF*-mutated disease as a distinct subtype within RAS pathway-aberrated leukemias. Drug sensitivity screens suggest broad CDK or HSP90 inhibition in addition to BRAF/RAS-directed inhibition may be effective targeted therapies in this prognostically poor AML subtype.

## Introduction

Mutations in the RAS/MAPK pathway genes, including *NRAS*, *KRAS*, and *PTPN11,* occur in >20% of patients with acute myeloid leukemia (AML), both *de novo* and secondary (1,2). Typically, these mutations are subclonal, suggesting they are later, transforming events in disease development (3). Mutations in *BRAF*, a gene encoding a protein kinase also in the RAS/MAPK signaling pathway, are well-known driver mutations in other cancer types (4,5), such as melanoma (6), papillary thyroid cancer (4), colorectal cancer (4) and hairy cell leukemia (7) . *BRAF* mutations are also recurrent mutational events in AML, albeit at a much lower frequency (3,8). While AML patients with *RAS* pathway mutations may respond to standard cytotoxic chemotherapy, depending on the molecular context, *BRAF* mutations in AML have been divergently associated with dismal outcomes (3). However, since many routinely used genetic testing panels are limited in their coverage of the *BRAF* gene, *BRAF* mutations might be overlooked and, their actual frequency, especially in relapsed/refractory disease, is likely underestimated. Importantly, in view of the prognostic impact of RAS pathway mutations on non-intensive treatment regimens, there is an urgent need to better understand the impact of *BRAF* mutations in AML, including their clinical, molecular, and cellular context, as well as their associations with response to intensive and non-intensive treatment regimens.

The most common *BRAF* mutation in cancer, known as p.V600E, substitutes the valine residue at codon 600 with glutamic acid (8). Some hematologic cancers, such as hairy cell leukemia, have recurrent *BRAF V600* mutations in almost 100% of cases (7). Notably, alternative non-canonical *BRAF* mutations exist in other cancer types and have been sorted separately from the V600E hotspot into two classes based on their kinase activity and oncogenic RAS*-*dependence (5). In total, three classes of *BRAF* mutations exist – class I (i.e. V600E) and class II mutations generate constitutively active kinases, but the class I mutations are unique in that these kinases can function as monomers (8–10). In contrast, class II mutant BRAF kinases still require dimerization for activity. Conversely, class III mutations are kinase-dead mutants with little to no BRAF kinase activity (11,12). However, class III mutant BRAF proteins are still able to amplify RAS pathway signals, but require an active or oncogenic RAS to transmit activation of the pathway. Little is currently known about the impact of the various classes of *BRAF* mutations on AML disease phenotype and therapeutic consequences, which may be attributed to the rarity of these mutations compared with other recurrent AML-associated mutations. However, the actual frequency of *BRAF* mutations is unknown, because most routinely used targeted next-generation sequencing panels cover only the V600 *BRAF* mutation.

In this study, we identified a cohort of *BRAF*-mutant AML patients and examined their leukemic molecular landscape, clonal architecture, and response to therapy. We found that *BRAF* mutations were highly enriched in myelodysplasia-related AML (AML-MR) and that most of them were non-V600E, non-canonical *BRAF* mutations, with each class of *BRAF* mutation showing a unique co-mutation and immunophenotype pattern. Moreover, we found that *BRAF*-mutant AML patients had dismal outcomes, and their disease showed resistance to venetoclax similar to *RAS*-mutated AML. However, through characterization of published *ex vivo* drug treatment data, we also identified unique drug sensitivities for *BRAF*-mutant AML that are not shared by other RAS-pathway mutant AML.

## Results

### Clinical analysis shows enrichment of BRAF mutations in AML-MR

We identified 50 patients with *BRAF* mutations (1% of all patients with AML treated at our centers during the study period) including 21 (42%) with newly diagnosed AML, 9 (18%) with relapsed/refractory AML, and 20 (40%) with newly diagnosed secondary AML, defined as having a documented antecedent chronic myeloid neoplasm or therapy-related AML. *BRAF*-mutant AML patients had a median age of 67 years (range, 19-84 yrs), and 54% of patients were male (**Supplementary Table 1**). The median variant allele frequency for all *BRAF* mutants was 15% (range, 1-83%). The median bone marrow blast percentage was 43% (range, 3-93%), and the median absolute monocyte count was 0.52/μL (range, 0-94.56). *BRAF* mutations were enriched in patients with AML-MR based on both the World Health Organization (WHO 5th) (13) (n=34/50, 68%) and the International Consensus Classification (ICC) (14) (n=26/50, 52%) classifications, including 11 (22%) patients who had a history of a previous MDS/MPN.

### Molecular Profiling of BRAF-mutant AML patients

The most frequent co-mutations in *BRAF*-mutant AML patients were in *TET2* (45%) and *ASXL1* (30%) (**Fig. 1A**). Oncogenic RAS pathway mutations were also common, with *NRAS* detected in 30% and *KRAS* in 23% of patients. In fact, 50% of *BRAF*-mutant patients (n=25/50) harbored at least one other oncogenic RAS/MAPK pathway mutation, suggesting RAS pathway addiction and that the RAS pathway in general may be a convergent mechanism of leukemogenesis for this AML subtype.

**Figure 1.**
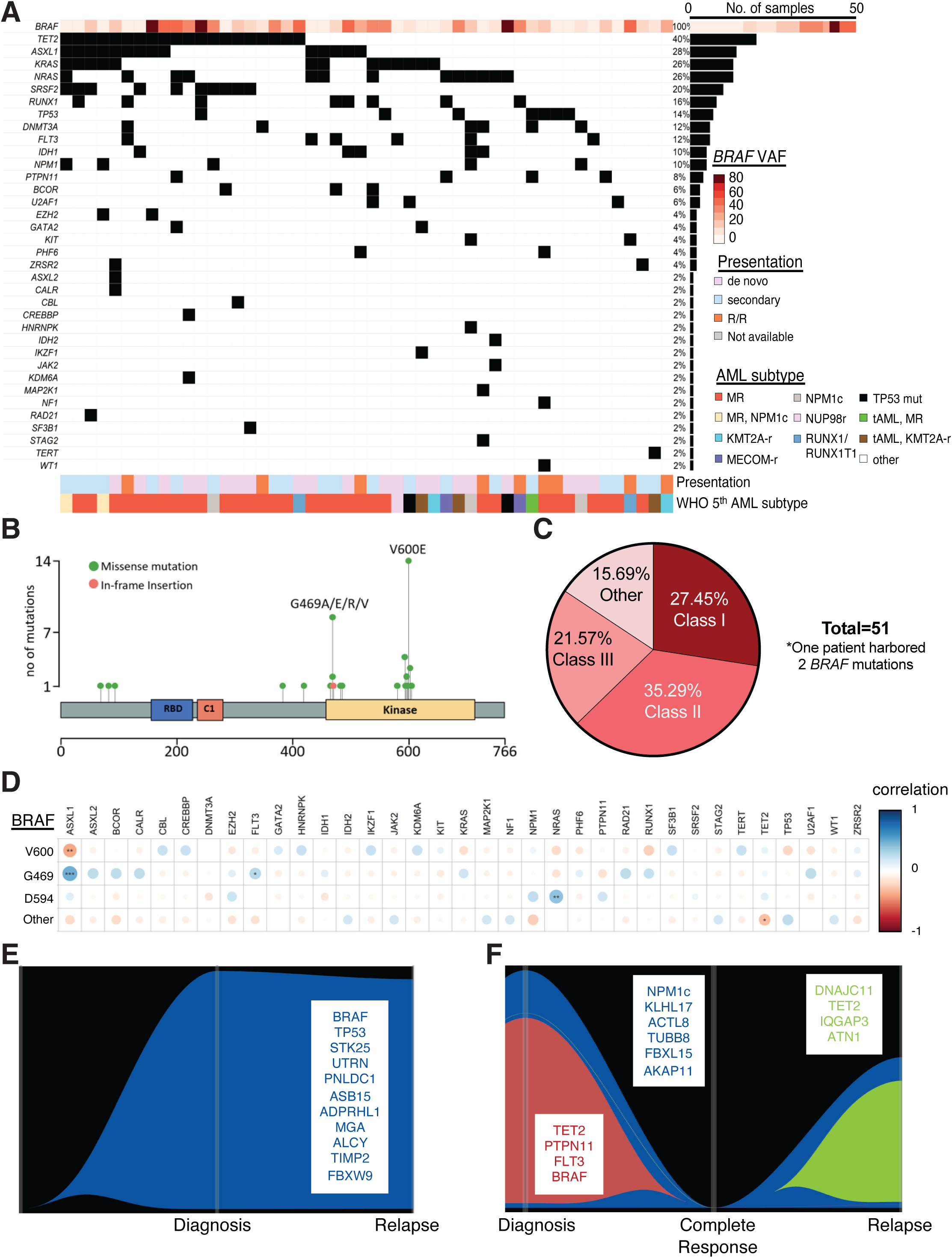
Non-canonical *BRAF* mutations in AML. **A)** Oncoprint of 50 *BRAF*-mutant AML patients and co-occurring mutations ranked by frequency in cohort (co-mutations present in ≥5% of cohort shown for clarity). AML type and disease presentation depicted on lower panel. **B)** Lollipop plot depicting frequency, gene location, and amino acid change of *BRAF* mutations identified in *BRAF*-mutant AML cohort. **C)** Pie graph depicting frequencies of canonical class I (V600E), non-canonical class II (including G469), and non-canonical class III (including D594) mutations. Mutations with no prior class distinction are denoted as other. **D)** Dot plot of enrichment (blue) or mutual exclusivity (brown) of AML co-mutations in each *BRAF* mutation class. Circles are colored and sized according to Pearson’s correlation coefficient, with *P*-values denoted as *P* <0.05, *; *P*<0.01, **; *P*<0.001, ***. **EF)** Fish plots of mutations identified from whole exome sequencing of two *BRAF*-mutant AML patients for whom paired relapse samples were available. Non-synonymous mutations identified are listed for each timepoint denoted by corresponding colors to identified clones.

Surprisingly, the majority (37/50, 73%) of *BRAF*-mutant cases were non-V600E variants (**Fig. 1B-C**). We found that 35% and 22% of patients harbored class II (n=18) and class III (n=11) mutations, respectively, with G469 (class II) and D594 (class III) mutations most common. This is in contrast to prior findings in solid tumors and lymphoid malignancies, where the V600E mutation represented the majority of *BRAF* alterations (4). As class II and class III mutations have different dependencies on RAS-pathway activation, we next examined co-mutations across the *BRAF* mutation classes identified in our cohort (**Fig. 1D**). We found that indeed patients with class III D594 mutations were significantly enriched with *NRAS* co-mutations (*P*=0.003). In contrast, class I V600 and class II G469 mutations seldom co-occurred with other RAS pathway mutations. Additionally, while *BRAF V600* mutations were mutually exclusive with *ASXL1* co-mutations (*P*=0.005), class II G469 mutations were significantly enriched in *ASXL1* co-mutated cases (*P*=0.0003). The *BRAF* class-specific co-mutational patterns with common myeloid malignancy mutations are suggestive of unique pathogenesis compared to other AML subtypes.

### Whole exome sequencing uncovers recurrent RAS-pathway addiction in BRAF-mutant AML

We performed whole-exome sequencing (WES) in a subset of patients (n=5) to identify mutations outside of the targeted sequencing panels. *BRAF*-mutant AML harbored a median mutational burden of 22 non-synonymous, coding mutations (range 13-32).

Interestingly, patients on average harbored 3 *RAS/MAPK* pathway mutations (range, 2-4; **Supplementary Table 2**). In two patients, WES was also performed at relapse. In one patient, the *BRAF* mutation was stable at relapse (**Fig. 1E**). In the second patient, the *BRAF* mutation was lost at relapse, but the patient instead acquired an *IQGAP3* mutation (**Fig. 1F**). IQGAP3 regulates cell proliferation by modulating the RAS/ERK signaling pathway, acting downstream of Rho family GTPases and interacting with ERK to promote cell cycle progression (15). These findings support a convergent reliance on the MAPK/RAS pathway in this leukemia subset.

### Single-cell molecular profiling uncovers unique co-mutation and immunophenotypic patterns

To resolve the clonal architecture of *BRAF*-mutant AML, we performed simultaneous single-cell DNA (scDNA) + cell surface protein analysis on 7 samples using the Mission Bio Tapestri platform (16,17). We found that *BRAF* mutations could be identified either in the dominant, largest non-wildtype clone (n=4) or in a sub-clone (n=3) (**Fig. 2A**). *BRAF* mutations co-occurred on a single-cell level with oncogenic *NRAS* and *KRAS* mutations but were mutually exclusive with a *PTPN11* mutation in one sample. This observation is significantly divergent from the clonal co-mutation patterns we previously observed in AML patients harboring signaling mutations (*RAS, FLT3*), where we very rarely observed more than one signaling mutation in a clone (16).

**Figure 2.**
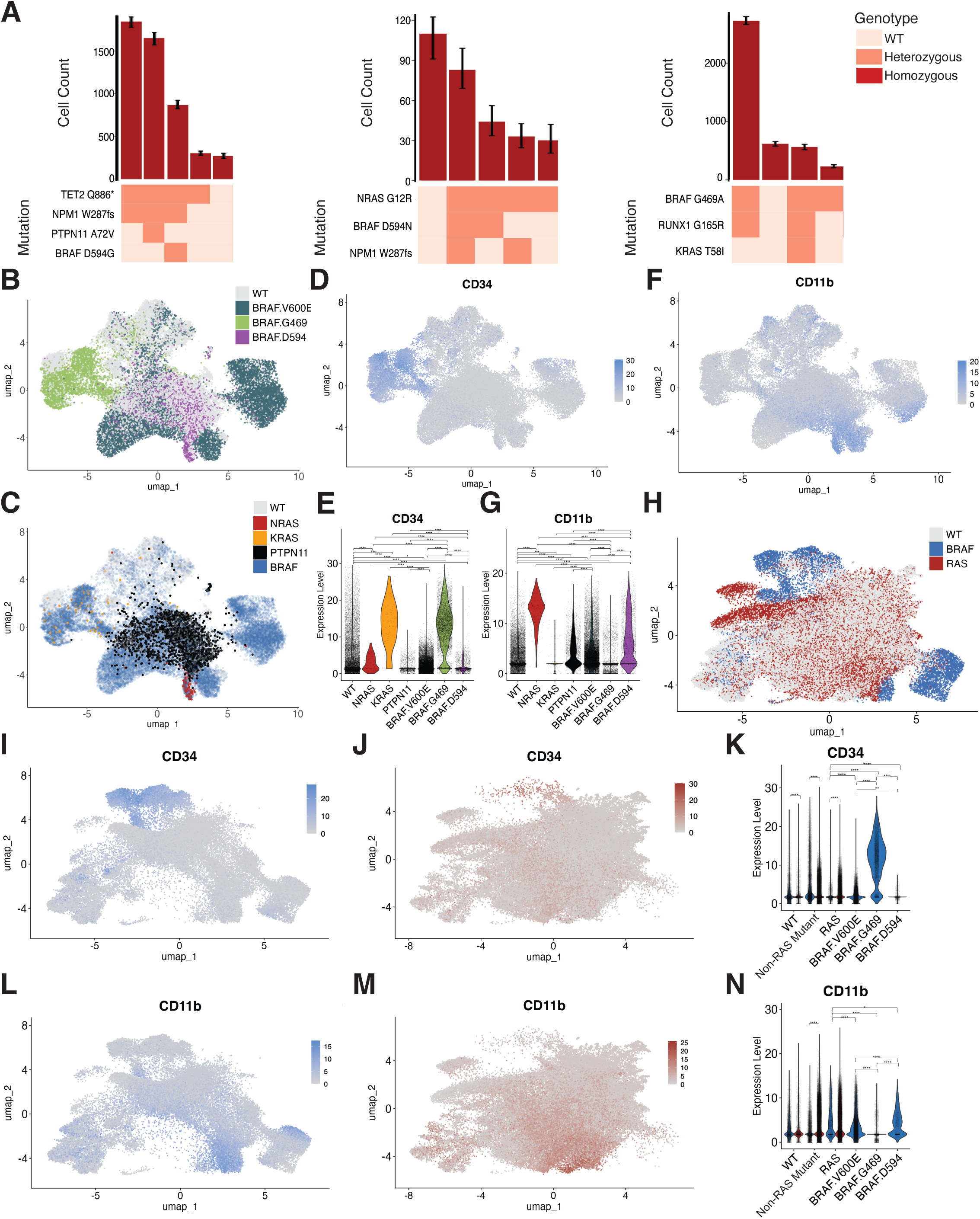
Genotype-immunophenotype relationships in *BRAF*-mutant AML clones. **A)** Clonographs from 3 representative *BRAF*-mutant AML patients. Clones were quantified and denoted in columns (upper bar plot) with genotype of the clone depicted in bottom heatmap. Zygosity of mutation in heatmap denoted as wild-type (WT; beige), heterozygous (orange), or homozygous (red). **BC)** UMAP of cells from *BRAF*-mutant AML patient samples (n=7) clustered by immunophenotype. Cells are denoted by (**B**) absence (grey) or presence of class I V600 (dark blue), class II G469 (green), or class III D594 (purple) *BRAF* mutations or (**C**) absence (grey) or presence of *NRAS* (red), *PTPN11* (black), *KRAS* (yellow), or *BRAF* (blue) mutations. **D-G)** UMAPs (**D, F**) and corresponding violin plots (**E,G**) of cells from A denoting low (grey) or high (purple) expression of CD34 (**D,F**) or CD11b (**E,G**) with colors in **F** and **G** denoted by genotype: WT (grey), *NRAS* (red), *PTPN11* (black), *KRAS* (yellow), *BRAF V600* (dark blue), *BRAF G469* (green), and *BRAF D594* (purple). Each dot within violin plot represents a cell. **H**) UMAPs of cells from A (n=7 samples) and *BRAF*-WT/*RAS*-mutant cells (n=6) clustered by immunophenotype. Cells are denoted by absence (grey) or presence of *BRAF* (blue) or other RAS pathway (red) mutations. **IJ)** Faceted UMAP from (**H**) denoting *BRAF*-mutant cells (**l**) and *BRAF-* WT*/RAS*-mutant cells (**J**) with low (grey) or high (red) expression of CD34. **K**) Corresponding violin plot for CD34 expression from cells in **IJ**. **LM**) Faceted UMAP from (**H**) denoting *BRAF*-mutant cells (**L**) and *BRAF-*WT*/RAS*-mutant cells (**M**) with low (grey) or high (red) expression of CD11b. **N**) Corresponding violin plot for CD11b expression from cells in **LM**. Asterisks denote *P* <0.05, *; *P*<0.01, **; *P*<0.001, ***, *P*<0.0001, **** (**EGKN**).

We next investigated immunophenotype patterns in *BRAF*-mutant AML samples. We found that *BRAF* mutations were present in cells spanning different lineage states, highly dependent on the type of *BRAF* mutation (**Fig. 2B**). Similarly, we found that other RAS pathway co-mutations in *BRAF*-mutant samples spanned different lineage states (**Fig. 2C**). Specifically, we found *BRAF G469* mutant clones to be significantly enriched for a stem/progenitor phenotype with CD34 expression (*P*<2x10^-16^) (**Fig. 2D-E)**. *KRAS*-mutant cells were also significantly enriched in expressing CD34 in *BRAF*-mutant samples (*P*<2x10^-16^). Conversely, *BRAF V600E* and *BRAF D594* mutant clones were enriched in monocytic cells expressing CD16 and/or CD11b (*P*<2x10^-16^ for all) (**Fig. 2F-G, Supplementary Fig. S1A**). *NRAS* and *PTPN11* mutant clones were also found to have increased CD11b and CD16 expression, as previously identified (16–18) (**Fig. 2E-G, Supplementary Fig. S1A-B**). Compared to BRAF WT clones, all *BRAF*-mutant clones had higher CD123 expression (**Supplementary Fig. S1C-D**). *BRAF V600E* mutant clones, specifically, displayed higher expression of FLT3 compared to class II and class III mutant clones (**Supplementary Fig. S1E-F**). These findings suggest that *BRAF* mutations from different classes may lead to highly divergent cell states and immunophenotypes of the resulting AMLs.

We then compared our *BRAF*-mutant AML samples to a separate subset of *BRAF* WT/*RAS*-mutant AML samples previously analyzed by scDNA+Protein (19), which harbor similar AML co-mutations found in a subset of our *BRAF*-mutant AML patients, including *NRAS* mutations (**Fig. 2H**). We found that *BRAF G469* mutant cells had overall higher expression of CD34 compared with *NRAS*-mutant cells from both sample cohorts (*P*<2x10^-16^; **Fig. 2I-K**, **Supplementary Fig. S1G**). Interestingly, even though *BRAF V600* and *BRAF D594* mutant clones expressed CD11b and CD14, the *NRAS*-mutant cells from *BRAF WT/RAS*-mutant samples expressed significantly higher levels of the monocyte markers (*NRAS* vs *BRAF V600*, *P*<2x10^-16^; *NRAS* vs *BRAF D594 P*=0.039) (**Fig. 2L-N, Supplementary S1G-J**). *BRAF V600* mutant clones had expression of CD123 comparable to that of *RAS* mutant clones from *BRAF WT/RAS*-mutant AML samples (**Supplementary Fig. S1G, S1K-M**).

*BRAF mutations are associated with poor overall survival regardless of treatment regimen* The overall response rate (CR, CRi, or CRh) was 46% and the median OS was 5.67 months (range, 0.13-90.7 months) for the entire cohort (**Supplementary Fig. S2A**). Newly diagnosed (ND) *BRAF*-mutant patients were treated with a variety of low-intensity (LI, n=7) and high-intensity (HI, n=14) treatment regimens with or without venetoclax (20) (**Fig. 3A**). ND *BRAF*-mutant AML patients had poor overall survival (OS) regardless of therapeutic regimen (*P* = 0.2752; LI+/-VEN: 15.09 months, 95% Cl 0.13-non-estimable (NE) vs HI+/-VEN: 13.90 months, 95% Cl 2.37-NE). Surprisingly, *BRAF* variant allele burden did not have a significant impact on OS in newly diagnosed patients (VAF≥15%: 5.02 months, 95% CI: 0.43 to 22.68 vs VAF<15%: 18.47 months, 95% CI: 1.02 to NE; P = 0.132; **Fig. 3B**). Prior exposure to venetoclax among R/R AML did not impact OS (**Supplementary Fig. S2B**, *P* = 0.884). We next reviewed the BeatAML cohort (2,21) and identified three ND patients with *BRAF* mutations. Their OS was very short, with all three patients deceased at 4 months post-study enrollment. Interestingly, patients with *BRAF*-mutant AML from the BeatAML cohort had significantly higher percentages of monocytes (P < 0.0018) in the peripheral blood compared to *NRAS/KRAS*-mutant patients (**Fig. 3C**).

**Figure 3.**
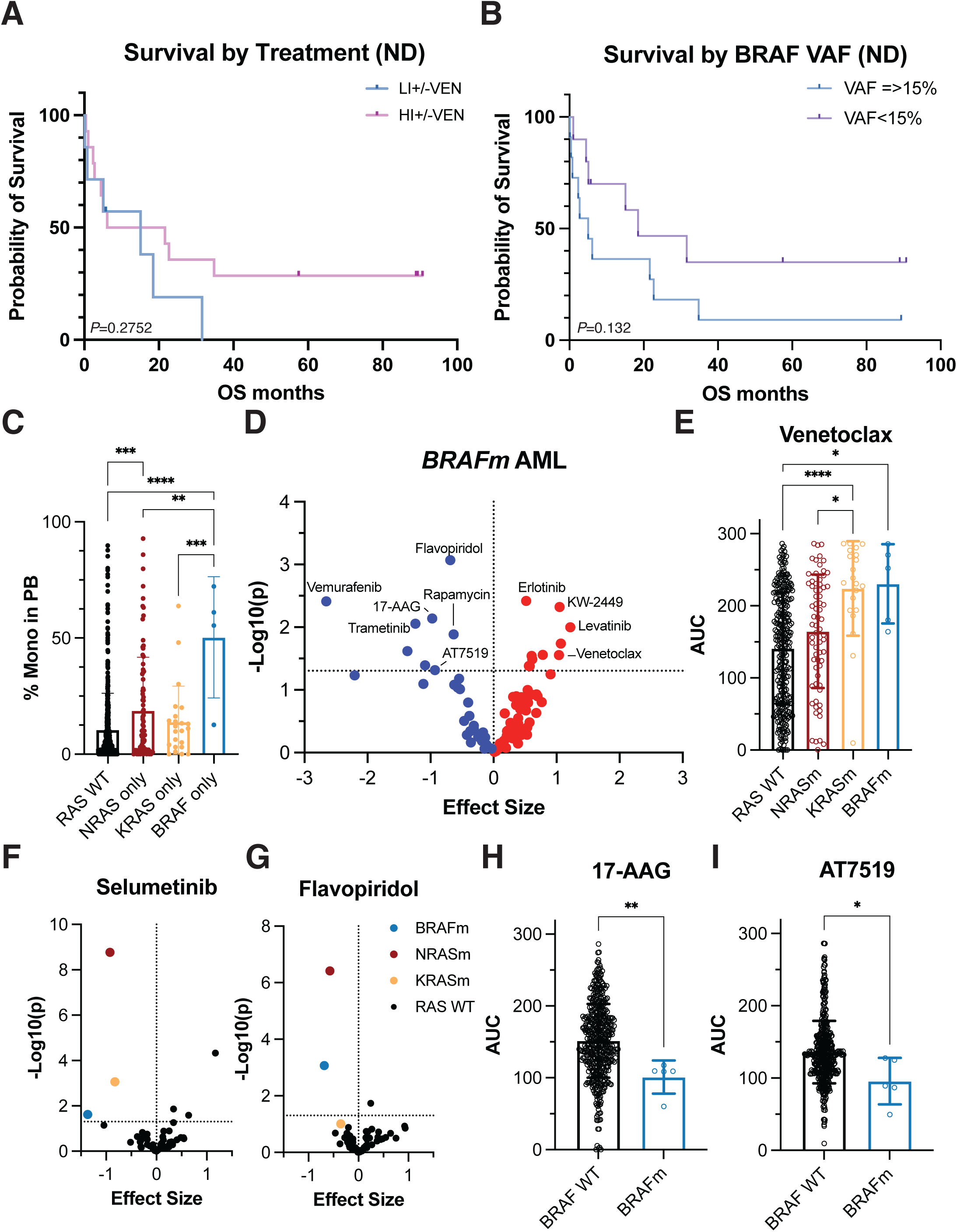
*BRAF*-mutant AML patients have significantly poor overall survival and unique drug sensitivities. **A)** Kaplan-Meyer survival curve of *BRAF*-mutant patients treated with low-intensity chemo +/-venetoclax (LI+/-VEN; blue) or high-intensity chemo +/-venetoclax (HI+VEN; pink). *P*-value calculated by logrank test. **B)** Kaplan-Meyer survival curve of *BRAF*-mutant patients stratified by *BRAF* mutant variant allele frequency (VAF). Patients with a VAF greater than or equal to 15 (n = 25, blue) are compared to patients with a VAF less than 15 (n=25, purple). *P*-value calculated by logrank test. **C**) Violin plot showing percentages of monocytes in the peripheral blood for BeatAML cohort newly diagnosed AML patients harboring *BRAF* (n=3; blue), *NRAS* (n=64; red), and *KRAS* (n=22; yellow) mutations compared to *RAS-WT* AML patients (n=433; black). **D)** Volcano plot showing increased sensitivity (blue) or resistance (red) of primary AML samples harboring *BRAF* mutations (both V600E and non-canonical mutations) to short term *ex vivo* drug treatments by plotting Glass’ Delta effect size compared to all other tested AML samples (*BRAF*-WT). Dotted line indicates *P* value = 0.05. Data from BeatAML cohort. Specific drugs discussed in the text are highlighted. **E)** Bar plot denoting area under the curve (AUC) results for RAS WT (black, n=295), NRAS-mutant (red; n=69), KRAS-mutant (yellow; n=21), or BRAF-mutant (blue; n=5) samples treated with venetoclax. **FG)** Volcano plots of the relative sensitivity/resistance of *BRAF* (blue), *NRAS* (red), and *KRAS* (yellow) mutant AML samples to Selumetinib (**F**), and Flavopiridol (**G**) compared to samples from all other *RAS WT* AML genotypes (black). Dotted line indicates *P* value = 0.05. **HI)** Bar plots depicting AUC results for *BRAF-*WT (black) and *BRAF*-mutant (blue) samples treated with 17-AAG (**H**) or AT7519 (**I**). Asterisks denote *P* <0.05, *; *P*<0.01, **; *P*<0.001, ***, *P*<0.0001, **** (**DFHI**).

### Ex vivo drug sensitivity data suggest BRAF-specific vulnerabilities

To further evaluate treatment options in *BRAF*-mutant AML, we looked at relative drug efficacies using the samples in the BeatAML cohort (2,21). Five *BRAF*-mutant AML samples were tested, including 3 harboring V600E mutations and 2 harboring class II G469 mutations (**Fig. 3D**). *BRAF*-mutant AML samples were significantly more resistant to 8 therapeutic agents compared with *BRAF*-WT AML samples. These drugs included Erlotinib, an EGFR inhibitor, and two multi-kinase inhibitors, KW-2449 and Lenvatinib. Interestingly, we also found that *BRAF*-mutant AML samples showed increased resistance to venetoclax compared to RAS-WT patients, providing a rationale for the poor responses to venetoclax-based regimens seen in our patients (*P* = 0.049; **Fig. 3AE, Supplementary Fig. S2B**). *NRAS/KRAS*-mutant AMLs have been previously shown to be resistant to venetoclax-based regimens and showed resistance levels similar to *BRAF*-mutant AML (18,22–24). Conversely, these samples were found to have significantly increased sensitivity to 8 therapeutic agents compared to *BRAF*-WT AML, with the highest sensitivity to flavopiridol, a cyclin dependent kinase (CDK) inhibitor (25), which had previously shown exciting efficacy data in specific AML subtypes, and had been more recently suggested as a strategy to overcome venetoclax resistance(26). Not surprisingly, additional efficacies were observed upon treatment with vemurafenib, a mutant BRAF V600E inhibitor (27), trametinib, a MEK inhibitor (28), and rapamycin, an mTOR complex inhibitor(29), and also with 17-AAG, an HSP90 inhibitor (30).

We compared the drug efficacies of *BRAF*-mutant AML to those identified in *NRAS/KRAS*-mutant AML samples. *RAS*- and *BRAF*-mutant samples all shared sensitivities to Trametinib, Selumetinib, and CI-1040, compared to other AML genotypes (**Fig. 3F, Supplementary Fig. S2C**). *NRAS-* and *BRAF-*mutant AML shared sensitivity to flavopiridol, but *KRAS*-mutant AMLs were not significantly sensitive (**Fig. 3G**). Conversely, *NRAS*-mutant AML was sensitive to panobinostat; however, the responses of *KRAS*- and *BRAF*-mutant AML did not reach statistical significance (**Supplementary Fig. S2D**). *BRAF*-mutant AML showed unique sensitivity to the HSP90 inhibitor, 17-AAG and the broad CDK inhibitor, AT7519, compared to *BRAF WT* AMLs and trended towards significance with *RAS*-mutant AMLs (**Fig. 3HI, Supplementary Fig. S2E**). These findings suggest that *BRAF*-mutant AML may have unique drug sensitivities that extend beyond BRAF/RAF pathway inhibition, with specific promise of both HSP90 and broad CDK inhibition that should be studied further as alternative therapeutic options for patients.

## Discussion

Oncogenic mutations in the RAS/MAPK signaling cascade occur in almost all types of human cancer. Alterations in the BRAF gene occur in approximately 15% of all human cancers (4). In AML, *RAS/MAPK* pathway mutations occur in approximately 20% of patients, most commonly in *NRAS* (1,3). Interestingly, while *RAS* pathway mutations are frequent in AML, mutations in the gene encoding the downstream effector kinase *BRAF* occur much less frequently. This is in contrast to many solid malignancies where *BRAF* mutations represent significant cancer drivers, and to the B-cell derived hairy cell leukemia, where *BRAF* V600 mutations occur in almost 100% of cases. However, almost 75% of the *BRAF* mutations in our patient cohort were non-canonical mutations that were not located at the V600 hotspot. Since *BRAF* V600E mutation makes up 70-80% of all *BRAF* gene alterations in human cancer, our findings suggest that AML is unique in being enriched for non-hotspot *BRAF* mutations.

BRAF mutant proteins have been classified based on the mutation location and the intrinsic kinase activity of the mutated protein. Class I (V600) and class II (i.e. G469 and others) mutants function as RAS-independent, hyperactive monomers and dimers, respectively (8,9). Class III (i.e. D594 and others) mutants, on the other hand, are inactive or kinase-dead mutants and require an activated or oncogenic RAS protein to perpetuate the signaling pathway (11,12). Indeed, *BRAF D594* mutant AML patients in our cohort frequently harbored *NRAS* co-mutations, and our single-cell data from a *BRAF D594* mutant patient showed that the *BRAF* and *NRAS* mutations co-occurred in the same clone, indicating the potential dependence of *BRAF D594* mutations on activated RAS in AML as well. Interestingly, we observed divergent co-mutation patterns between V600 and G469 mutations, particularly with *ASXL1* co-mutations, suggesting the *BRAF* mutation classes may divergently synergize with recurrent AML mutations. Similarly, we also found unique immunophenotype patterns between the different *BRAF* mutant clones, with *BRAF G469* mutant clones enriched in CD34 expression and *BRAF D594* mutant clones having higher expression of CD11b and CD16. Additionally, our *BRAF*-mutant AML samples showed higher levels of CD123 expression, which warrants testing the efficacy of recent CD123-targeted therapies (31). Not only do these findings suggest that the *BRAF* mutations may be prevalent with certain co-mutations based on synergistic interactions, but also that *BRAF* mutant classes are enriched in distinct leukemic cell compartments. Although limited by the number of samples available, the results of our multiomic analysis underscore the divergent phenotypes among AML samples with different *BRAF* mutations and suggest that further studies are needed to determine how the *BRAF* mutations interact with recurrent AML drivers and affect the phenotype of the leukemia.

While *RAS* mutations in other cancers typically bestow poor patient outcomes, the prognostic relevance of *RAS* mutations remains less clear in AML. *NRAS*-mutant AML patients typically show superior responses to standard chemotherapy compared with patients with other signaling mutations (3), but *RAS/MAPK* pathway mutations are also enriched in AML patients who are either refractory to or who develop resistance to targeted and novel therapies. *BRAF* mutations in AML have been associated with poor prognoses; however, many studies lacked sufficient numbers of *BRAF*-mutated cases for analysis. Our study shows poor outcomes of *BRAF*-mutated AML patients, regardless of the frontline regimen. These findings align with the recurrent acquired resistance/relapse mechanism for the venetoclax/azacitidine combination treatment, which lies in the expansion of a myelomonocytic blast population enriched for *NRAS/MAPK* mutations(18,23). This is further supported by our drug sensitivity data on *BRAF*-mutant AML obtained from the BeatAML cohort, which suggests that cells harboring *BRAF* mutations are more resistant to venetoclax; however, it is still unknown if they utilize similar mechanisms as *N/KRAS*-mutant cells. These findings suggest that *BRAF* mutations, like other *RAS/MAPK* pathway mutations, appear to play a significant role in therapy resistance and may serve as a convergent mechanism for therapy escape in AML.

Studies in other cancer subtypes have uncovered that each class of *BRAF* mutant proteins has distinct biochemical and signaling properties, which may lead to different dependencies on interacting proteins and alternative pathways. For example, these studies suggest that class III BRAF mutants are sensitive to RAS-pathway inhibition (12), whereas the class II BRAF mutants may be sensitive to MAPK-pathway inhibitors outside of RAS-targeted compounds (32). Data from the BeatAML study uncovered multiple therapeutic agents that may hold promise in *BRAF*-mutant AML patients, including HSP90 and CDK inhibition. However, the majority of the 5 samples harbored the V600E mutation, and no samples tested contained a class III mutation. Therefore, further examination of these agents in class II and III mutant AML samples is required to fully characterize their efficacy as potential avenues for *BRAF*-mutant AML treatment.

In conclusion, our study demonstrates that while rare, *BRAF* mutations are recurrent in AML, and are typically non-canonical. They exhibit distinct, class-specific co-mutation and immunophenotypic patterns, and are associated with poor survival outcomes, even in newly diagnosed patients. *Ex vivo* drug sensitivity studies identified potential therapeutic agents for *BRAF*-mutant AML. These findings highlight the unique nature of *BRAF*- mutated AML, underscoring the critical need for improved understanding of this genotype to enhance survival outcomes, and the inclusion of expanded *BRAF* gene coverage into routine testing panels for newly diagnosed and relapsed/refractory AML patients.

## Methods

### Patient Cohort

This study was approved by CCHMC IRB (protocol 2022-0806, for multiomic profiling of de-identified samples), OSU IRB (protocol 2023C0062), MDACC IRB (protocol PA17- 0485), and OHSU IRB (protocols 9570; 4422, for ex vivo drug assays and multiomic profiling). Patient consent was obtained according to protocols approved by the institutional IRBs in accordance with the Declaration of Helsinki. Patients treated on Alliance for Clinical Trials in Oncology (Alliance) protocols gave informed consent for tissue banking and sequencing under CALGB 9665 and 20202. No clinical and/or outcome data analyses were performed on these patients. Diagnosis according to WHO classification (13) and ICC (14) criteria and disease status assignment are provided in Supplementary Table 1. Mononuclear cells were isolated via Ficoll gradient centrifugation from obtained bone marrow aspirates or peripheral blood specimens. Single-cell suspensions of samples were viably frozen and stored in liquid nitrogen.

### Bulk DNA Molecular Profiling

All patients and corresponding samples included in the BRAF-mutant clinical cohort and control cohorts underwent bulk DNA molecular profiling. Samples in the BeatAML cohort obtained from OHSU were sequenced with a combination of whole exome sequencing, targeted sequencing panels, and clinical genetic platofrms (Sequenom and GeneTrails (2,21). Analysis of samples collected at MDACC used targeted PCR-based NGS panels covering 81 recurrently mutated genes (33) or 28 recurrently mutated genes (34) in myeloid disorders. The Alliance samples were sequenced using an NGS panel covering 80 cancer and/or leukemia-associated genes, as described previously(35). OSU patient sequencing was obtained from the patients’ clinical records using the OSU CLIA-approved panels.

### Whole Exome Sequencing

Whole-exome hybrid capture and NGS sequencing were performed on a subset of BRAF-mutant AML samples in our study at OSU. The capture included chromosomal tiling probes (250-fold coverage) and was performed on leukemic and matched germline tissue for each patient for the detection of single nucleotide variants and small insertions/deletions, as described previously (36).

### Reagents

Mission Bio Tapestri-related reagents were included in the Myeloid DNA+Protein V2 sequencing kit purchased from the company with the following exceptions: Cell Staining Buffer, TotalSeqD CD135 antibody, and TotalSeqD Antibody Cocktail v2 were purchased from Biolegend. The Myeloid amplicon panel is a previously described (37), commercially available panel from Mission Bio, which targets 45 genes via 312 amplicons.

### Single-Cell Multi-omic Sequencing

Samples from AML patients were thawed, washed with FACS buffer (PBS supplemented with 2% fetal bovine serum), filtered to obtain single-cell suspensions, and quantified using a CellDrop (Denovix). Cells (1x10^6^ viable cells) were then incubated with Tapestri blocking buffer and TruStainFcX (Biolegend) on ice for 15min followed by a 30min incubation with the TotalSeqD Antibody Cocktail supplemented with 2μL of TotalSeqD CD135 on ice. Stained cells were then further processed as previously described in commercially available protocols(16,19,38). Sequencing of pooled libraries were performed by the DNA Genomic Sequencing shared facility at CCHMC.

### Single-cell DNA Sequencing Analysis

Sequencing reads were processed as follows: reads trimmed, aligned to the hg19 human genome, assigned barcodes, and genotypes were called using GATK by the cloud-based Mission Bio Tapestri v2 pipeline. Further analysis of processed H5 files was performed using the scDNA package (https://github.com/bowmanr/scDNA, v1.01) in R v4.3. In the scDNA package, H5 files from the Tapestri pipeline were input and variants of interest were identified based on clinical sequencing identified variants. All variants included were manually investigated in IGV. We selected non-synonymous, exonic variants that were genotyped in >50% of assayed cells and had a computed VAF >1%. The variant list was further refined to exclude variants that were 1) confirmed SNPs, 2) recurrently mutated broadly across cohorts at a fixed VAF, and/or 3) represented exclusively in low-quality reads or clipped reads. The ‘tapestri_h5_to_sce’ function from the scDNA package was used after variant selection to generate a SingleCellExperiment class object using the default cutoffs of allele frequency (AF) variance >25, depth (DP) >10, and genotype quality (GQ) >30. The AF variance refers to the maximum deviation from 50% by which a heterozygous call should be masked as inaccurate. We only retained variants that passed all three filters in over 80% of cells and only cells that passed all three filters were included in the final analysis and termed “Complete” cells, indicating they received a reliable genotype for all genes of interest. Clones were identified and statistically summarized following variant identification using the ‘enumerate_clones’ and ‘compute_clone_statistics’ functions, respectively.

### Single cell DNA+Protein (scDNA+Protein) Sequencing Analysis

Protein matrices were extracted from Tapestri pipeline generated H5 files using the scDNA package after genotyping and clone enumeration above. The SingleCellExperiment object was converted to a Seurat object (v5.1) with metadata containing the genotyping information. All complete cells identified above were bound to a single protein matrix for cohort level protein analysis across all samples. Protein data was normalized across cells using DSB, scaled across all samples, and analyzed by PCA. Samples were integrated with Harmony, clustered (SLM) and then subsequently visualized by Uniform Manifold Approximation and Projection (UMAP) (39). Cell type calls were performed by manual interpretation of protein expression. Data were visualized using Seurat, ggplot2, gridExtra, ggpubr, magick, patchwork, raster and scCustomize packages (https://samuel-marsh.github.io/scCustomize/).

### Beat AML Ex Vivo Drug Screen Data

The *ex vivo* functional drug screens were performed as described previously (2,21) on freshly isolated mononuclear cells harvested as part of the BeatAML cohort. Data shown in Figure 3 and Supplementary Figure 2 were generated as part of the BeatAML cohort and data downloaded from their online browser (http://vizome.org/aml2). Data points were replotted using GraphPad Prism v10.

### Statistical Analysis

Wilcoxon rank tests were used to assess significant differences in protein expression in the scDNA+Protein data (**Fig. 2, Supplementary Fig. S1**). Survival data were analyzed by logrank (Mantel-Cox) tests (**Fig. 3, Supplementary Fig. S2**). Comparisons of area under the curve (AUC) metrics from the BeatAML *ex vivo* drug data were analyzed by Kruskal-Wallis tests (**Fig. 3D-E, Supplementary Fig. S2D-E**) or Mann-Whitney tests (**Fig. 3H-I**) when appropriate.

### Plotting and Graphical Representations

The oncoprint in Fig. 1A was generated using oncoPrint package in R. Clonographs and UMAPs from the single-cell multiomic profiling were generated in R. Graphs unless otherwise noted were generated using GraphPad Prism v10. UMAP data was plotted using the ggplot2 package (RRID:SCR_014601) in R. Other data processing was performed in R utilizing packages including: tidyr, dplyr, RColorbrewer, pals, and cowplot.

## Data and Code Availability

All scripts and processed data files are available at https://github.com/bowmanr/scDNA and https://github.com/plask3189/scProtein_Analysis for DNA+Protein analyses. Raw data files are available upon request from the authors and are being uploaded to dbGAP prior to final publication.

## Acknowledgements

We thank the CCHMC Genomics Sequencing Core (RRID:SCR_022630) for performing library sequencing. L.A.M. is supported by a National Cancer Institute grant (R00 CA252005) and an American Society of Hematology (ASH) Junior Faculty Scholar award.

R.L.B. is supported by a National Cancer Institute grant (R00 CA248460) and an ASH Junior Faculty Scholar award. This work was also supported by the ASH Junior Faculty Scholar award, the NCI R00 award, an American Cancer Society Institutional Research Grant award, and a University of Cincinnati Cancer Center Pilot Award to L.A.M. This work was also supported by a summer research award from the Carey & Stella Wamsley Charitable Trust (through University of Cincinnati College of Medicine Office of Research) to D.L. The authors are grateful to the patients who consented to participate in these clinical trials and the families who supported them; to Christopher Manring and the CALGB/Alliance Leukemia Tissue Bank at The Ohio State University Comprehensive Cancer Center, Columbus, OH for sample processing and storage services; and to Lisa J. Sterling for data management.

## Author Contributions

S.L., A-K.E., and L.A.M. conceptualized studies. L.A.M. and R.L.B. designed and optimized single-cell DNA/Protein experimental methodologies and bioinformatic workflow. Y.A-S., D.N., K.M., M.J.R., K.P.P., C.J.W., J.B., A.R.S., A.L., C.D.D., N.G.D., T.M.K., F.R., A.J.C., J.E.K, B.L.P., W.G.B., M.R.B., G.M., G.L.U, W.S., R.M.S., L.J.M., J.C.B., J.S.B., J.T., S.L., and A.-K.E provided de-identified patient samples, annotated clinical information, and performed analysis of clinical data. D.L. performed scDNA+Protein library preparation and sequencing. K.P. performed all computational multiomic analysis with guidance from R.L.B. J.T. provided ex vivo drug sensitivity data from BeatAML cohort. D.L., K.P., S.L., A.-K., E. and L.A.M. generated manuscript figures. S.L., A.-K. E. and L.A.M. funded the study. K.M., S.L., A-K.E. and L.A.M. wrote and edited the manuscript. All authors read the manuscript and agreed on the final version.

## Conflicts of Interest

L.A.M. and R.L.B. had previously received honoraria for speaking arrangements and had previously served on a Speakers Bureau for Mission Bio, Inc. J.C.B. has ownership interest in Vincera, an advisory and consultancy role with Novartis, Syndax, and Vincera, research funding from Genentech, Janssen, Acerta, and Pharmacyclics, an AbbVie company. A.-K.E. has received an honorarium from AstraZeneca for serving on their Diversity, Equity, and Inclusion Advisory Board and has received a research grant from Novartis. Spouse of A.-K.E. has ownership interest in Karyopharm Therapeutics. SL has research funding from Amgen and Astellas and has received honoraria/consultation fees from Abbvie, Arima, BMS, Caris, Cogent, Daiichi Sankyo, Johnson and Johnson, Kura Oncology, Qiagen, Recordati, Servier, Stemline Therapeutics, Syndax, Tempus AI. The other authors declare no competing interests.

---

**Supplementary Fig. S1.**
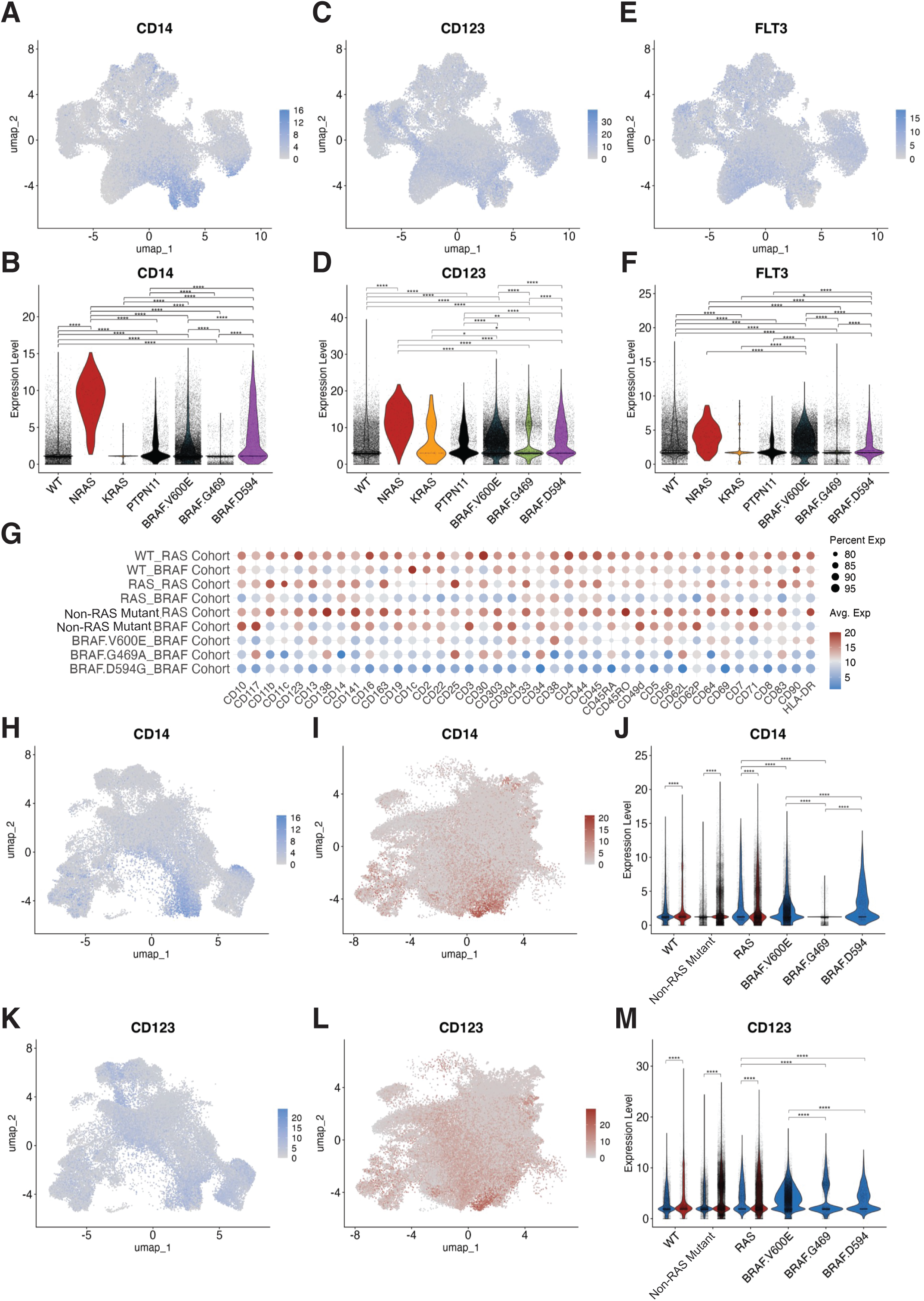
A-F) UMAP (A,C,E) and corresponding violin plots (B,D,F,) of cells from Fig. 2A denoting low (grey) or high (purple) expression of CD14 (**AB**), CD123 (**CD**) or FLT3 (**EF**) with colors in **B**, **D**, and **F** denoted by genotype: WT (grey), *NRAS* (red), *PTPN11* (black), *KRAS* (yellow), *BRAF V600* (dark blue), *BRAF G469* (green), and *BRAF D594* (purple). Each dot within violin plot represents a cell. **G**) Dot plot of row normalized expression of each cell surface protein marker for clones from *BRAF*-mutant samples and clones from *BRAF-*WT*/ RAS*-mutant samples. **HI**) Faceted UMAP from (Fig. 2H) denoting *BRAF*-mutant cells (**H**) and *BRAF-*WT*/RAS*-mutant cells (**I**) with low (grey) or high (red) expression of CD14. **J**) Corresponding violin plot for CD14 expression from cells in **HI**. **KL**) Faceted UMAP from (Fig. 2H) denoting *BRAF*-mutant cells (**K**) and *BRAF-* WT*/RAS*-mutant cells (**L**) with low (grey) or high (red) expression of CD123. **M**) Corresponding violin plot for CD123 expression from cells in **KL.** Asterisks denote *P* <0.05, *; *P*<0.01, **; *P*<0.001, ***, *P*<0.0001, **** (**BDFJM**).

**Supplementary Fig. S2.**
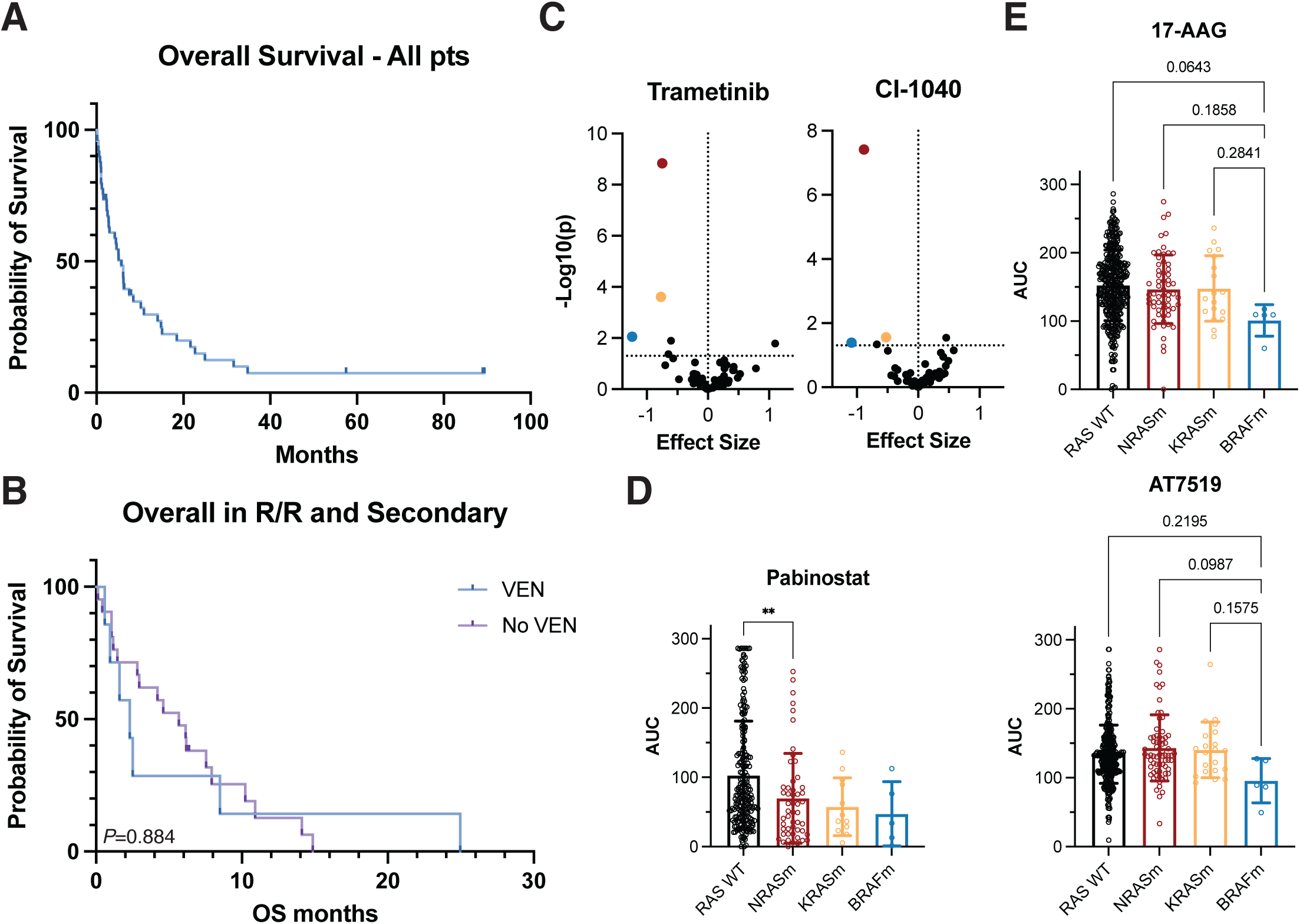
AB) Kaplan-Meyer survival curves of (**A**) all *BRAF*-mutant patients (n=50), and (**B**) relapse/refractory and secondary AML *BRAF*-mutant patients (n=28) stratified by whether prior treatment regimens had included venetoclax (n=7, blue) compared to not (n=21 purple). **C)** Volcano plots of the relative sensitivity/resistance of *BRAF* (blue), *NRAS* (red), and *KRAS* (yellow) mutant AML samples to Trametinib (left) and CI-1040 (right) compared to samples from all other *RAS* WT AML genotypes (black). **DE**) Bar plots denoting area under the curve (AUC) results for *RAS WT* (black), *NRAS*- mutant (red), *KRAS*-mutant (yellow), or *BRAF*-mutant (blue) samples treated with (**D**) Pabinostat (WT: n=227, NRAS: n=52 KRAS: n=12, BRAF: n=5) or (**E**) 17-AAG (top, WT :n=402, NRAS: n=61, KRAS: n=18, BRAF: n=5), or AT7519 (bottom, WT: n=367, NRAS: n=70, KRAS: n=21; BRAF: n=5). *P* values provided or asterisks denote *P* <0.05, *; *P*<0.01, **; *P*<0.001, ***, *P*<0.0001, **** (**DE**).

**Supplementary Table 1.** Summary of pretreatment characteristics of *BRAF*-mutated patients with AML in our cohort.^a^

| <u>Demographic/Clinical Parameter</u> | <u>*BRAF*-mutant cohort</u> |
| --- | --- |
| Age at original diagnosis, years, median (range) | 67 (19-84) |
| <65 (%) | 23 (46) |
| ≥65 (%) | 27 (54) |
| Sex, number (%) |  |
| Female | 23 (46) |
| Male | 27 (54) |
| Disease at presentation, number (%) |  |
| De novo, newly diagnosed | 21 (42) |
| Relapse/refractory | 9 (18) |
| Secondary | 20 (40) |
| WHO Classification, number (%) |  |
| AML-MR | 34 (68) |
| AML with *NPM1* mutation | 3 (6) |
| AML with *MECOM*-rearrangement | 2 (4) |
| AML with *RUNX1:RUNX1T1* fusion | 2 (4) |
| t-AML with *KMT2A*-rearrangement | 3 (6) |
| Other | 6 (12) |
| ICC Classification, number (%) |  |
| AML-MR | 26 (52) |
| AML with mutated *NPM1* | 5 (10) |
| AML with mutated *TP53* | 6 (12) |
| AML-*KMT2A* | 5 (10) |
| AML with *MECOM*-rearrangements | 2 (4) |
| AML with *RUNX1:RUNX1T1* fusion | 2 (4) |
| Other | 4 (8) |
| Bone marrow blast percentage, % median (range) | 43 (3-93) |
| Absolute monocyte count, median (range) | 0.52 (0-94.56) |
| *BRAF* VAF, median (range) | 15 (1-83) |
| <15 (%) | 25 (50) |
| ≥15 (%) | 25 (50) |
| *BRAF* mutation class, number (%) |  |
| Class I | 13 (25.5) |
| Class II | 20 (39.2) |
| G469 | 13 (25.5) |
| Class III | 11 (21.6) |
| D594 | 7 (13.7) |
| Other | 7 (13.7) |

<sup>a</sup>Table provides median values for each parameter and number of patients and/or percentages of the cohort. For WHO and ICC classifications, classifications with more than 1 patient are shown with all others grouped in "Other".

**Supplementary Table 2.** Summary of whole exome sequencing (WES) results from 5 *BRAF*-mutant AML patients.^a-d^

| <u>Pt ID</u> | <u>Gene</u> | <u>Mutation Type</u> | <u>VAF Diagnosis</u> | <u>VAF Relapse</u> |
| --- | --- | --- | --- | --- |
| A | *TET2* | FS | 99.88 | NA |
|  | *ATP8B4* | SPL | 51.35 |  |
|  | *ECPAS* | SNV | 48.84 |  |
|  | *STOML2* | SNV | 48.83 |  |
|  | *NPM1* | FS | 48.00 |  |
|  | *NRAS* | SNV | 45.60 |  |
|  | *SRSF2* | SNV | 45.09 |  |
|  | *ABCG5* | SNV | 44.69 |  |
|  | *ARHGAP2* | SNV | 44.57 |  |
|  | *SHANK3* | SNV | 43.09 |  |
|  | *GRIP1* | SNV | 42.88 |  |
|  | ***BRAF*** | **SNV** | **42.28** |  |
|  | *MOV10* | SNV | 39.34 |  |
|  | *ZNF571* | FS | 34.27 |  |
|  | *ZNF850* | SNV | 11.60 |  |
|  | *AQP7* | SNV | 4.43 |  |
|  | *FAT3* | SNV | 4.31 |  |
|  | *RELN* | SPL | 3.25 |  |
| B | *NPM1* | FS | 60.58 | NA |
|  | *KAT6B* | FS | 57.16 |  |
|  | *ANKRD62* | SNV | 52.80 |  |
|  | *RNF213* | SNV | 50.86 |  |
|  | *KRAS* | SNV | 30.26 |  |
|  | *BICDL1* | SNV | 4.40 |  |
|  | *BOP1* | FS | 3.38 |  |
|  | *FAM104A* | FS | 3.19 |  |
|  | *DZIP1* | SNV | 3.05 |  |
|  | *CCDC50* | SPL | 2.90 |  |
|  | *RBM4* | DEL | 2.78 |  |
|  | *MRPS18A* | SNV | 2.19 |  |
|  | ***BRAF*** | **SNV** | **3.00** |  |
| C | *PRUNE2* | SNV | 50.50 | NA |
|  | *MCTP1* | SNV | 49.15 |  |
|  | *ESRP1* | SNV | 48.21 |  |
|  | *CDH23* | SNV | 46.52 |  |
|  | *KRAS* | SNV | 33.62 |  |
|  | *DCDC1* | TR | 25.83 |  |
|  | *DENND2A* | SNV | 22.95 |  |
|  | *NPY2R* | SNV | 18.91 |  |
|  | *ADAMTS1* | SNV | 9.63 |  |
|  | **BRAF** | **SNV** | **3.02** |  |
|  | SP8 | DEL | 2.99 |  |
|  | BMP2K | DEL | 2.44 |  |
|  | CNKSR2 | SNV | 2.26 |  |
|  | CCDC22 | SNV | 2.14 |  |
| D | DNAJC11 | SPL | 0.00 | 50.50 |
|  | NPM1 | FS | 51.81 | 50.00 |
|  | SCML1 | SNV | 46.30 | 49.76 |
|  | AKAP11 | SNV | 43.77 | 49.38 |
|  | NCKAP5 | SNV | 40.66 | 48.68 |
|  | FBXL15 | SNV | 50.15 | 47.46 |
|  | TET2* | TR;<br>SNV* | 50.47; 38.51;<br>0.32 | 46.96; 44.51;<br>0.0 |
|  | TUBB8 | SNV | 48.25 | 45.29 |
|  | ATP13A1 | SNV | 48.76 | 43.45 |
|  | ACTL8 | SNV | 51.89 | 43.24 |
|  | SIRT4 | SNV | 46.06 | 43.06 |
|  | GALK1 | SNV | 48.36 | 42.93 |
|  | LYN | SNV | 51.09 | 42.82 |
|  | KLHL17 | SNV | 43.21 | 42.09 |
|  | IQGAP3 | SNV | 0.07 | 42.06 |
|  | ADAMTS1 | SNV | 49.26 | 42.01 |
|  | NR1H2 | SNV | 40.62 | 40.96 |
|  | STAG2 | SNV | 36.86 | 40.87 |
|  | NYX | SNV | 40.71 | 40.86 |
|  | DMD | SNV | 49.58 | 39.69 |
|  | HNRNPM | SNV | 43.24 | 39.30 |
|  | C12orf4 | SNV | 49.27 | 39.28 |
|  | ATN1 | DEL | 0.00 | 33.08 |
|  | PUM1 | SNV | 0.00 | 3.54 |
|  | DNX35 | SPL | 0.00 | 3.10 |
|  | MAP4 | SPL | 0.39 | 2.79 |
|  | TRIP4 | SNV | 0.35 | 2.49 |
|  | SIGLEC16 | SNV | 0.94 | 2.46 |
|  | SOX1 | SNV | 1.32 | 2.19 |
|  | FLT3 | SNV | 22.73 | 0.00 |
|  | PTPN11 | SNV | 17.54 | 0.00 |
|  | **BRAF** | **SNV** | **4.55** | **0.00** |
|  | LAMB4 | SPL | 4.30 | 0.00 |
|  | WAC | FS | 3.59 | 0.00 |
| E | TP53 | SNV | 35.44 | 97.99 |
|  | LOC10192 | SNV | 50.07 | 53.01 |
|  | TMPRSS6 | SPL | 32.93 | 51.57 |
|  | FBXW9 | SNV | 39.27 | 49.92 |
|  | ASB15 | SNV | 51.79 | 48.88 |
|  | *CHST6* | FS | 51.01 | 47.87 |
|  | ***BRAF*** | **SNV** | **45.79** | **47.76** |
|  | *MROH2B* | SPL | 47.75 | 47.34 |
|  | *NRAS* | SNV | 42.12 | 45.90 |
|  | *NALCN* | SNV | 43.95 | 45.83 |
|  | *STK25* | SNV | 44.58 | 45.28 |
|  | *SLC4A7* | SNV | 42.97 | 45.20 |
|  | *ACLY* | SNV | 45.21 | 44.30 |
|  | *TIMP2* | SNV | 48.98 | 43.91 |
|  | *TJP1* | SNV | 52.52 | 42.68 |
|  | *PNLDC1* | SNV | 45.42 | 41.91 |
|  | *CHD1* | SNV | 42.46 | 41.60 |
|  | *UTRN* | SNV | 50.37 | 37.18 |
|  | *MGA* | SNV | 41.95 | 36.87 |
|  | *POGLUT3* | SPL | 0.00 | 10.66 |
|  | *VGLL4* | SNV | 0.47 | 4.71 |
|  | *ST6GAL1* | SPL | 0.00 | 4.38 |
|  | *SYNE1* | SNV | 0.00 | 4.37 |
|  | *DEAF1* | DEL | 3.17 | 4.32 |
|  | *RELN* | SNV | 0.00 | 3.15 |
|  | *CHRM4* | FS | 0.53 | 2.79 |
|  | *KLF2* | SNV | 0.00 | 2.52 |

<sup>a</sup>Patients A-C only had diagnosis sample available.
<sup>b</sup>Mutated genes listed in descending variant allele frequency (VAF) order in relapse sample or diagnosis sample if relapse sample not available (NA).
<sup>c</sup>Mutation types are denoted as FS: frameshift, SNV: single nucleotide variant, DEL: non-frameshift deletion, SPL: splice site mutation, and TR: truncation.
<sup>d</sup>Asterisk denotes more than 1 mutation or mutation type in same gene.

